## Supplementary Information for "Twist and Scout: Analysis and Curation of Particles in Cryo-Electron Tomography Using TANGO"

### 1. Subtomogram averaging of CA-SP1 capsid from VLPs of immature HIV-1

The number of particles was determined based on the lowest binning occupancy, which corresponded to the bin with mean angular score 0.0 - 0.2. This bin contained 126 particles. From other bins, 126 particles were selected randomly to even the number of particles in each category. First, only averages based on the original alignment from the full particle set were generated (Supplementary Figure 5 **a**) using four times binned subtomograms. One can observe deterioration of the hexagonal lattice with decreasing mean angular score – at the third bin (mean angular score 0.4 - 0.6), the lattice starts to be incomplete and disappears completely for a mean angular score of 0.0 - 0.2.

We proceeded with alignment for all five bins containing 126 particles. First, the in-plane angles were randomized and a new average was computed (Supplementary Figure 5 **b**), again using four times binned subtomograms. Already at this stage, one can observe differences between the different bins; while the average in the bin with the highest mean angular scores maintains the characteristic spherical curvature in the side views, this curvature disappears in the lower bins (below mean angular score of 0.8). The initial averages and randomized motive lists were used for subsequent alignment. In total, fifteen iterations were run without imposing any symmetry during alignment (see Supplementary Table 5 for parameters). The results are shown in Supplementary Figure 5 **c**. While particles from the bin with the highest mean angular score recovered the hexagonal lattice quite well, the lattice was only partially recovered in the second bin and further deteriorated in the bins with lower scores. STA was performed using novaSTA [50].

### 2. Additional descriptors

#### 2.1. PL descriptor

The two descriptors that stem from analyses of triangulated 2D surfaces can be helpful when investigating lattice symmetries, e.g. when computing how much the neighborhood of a vertex (specifically: the star of the vertex, Supplementary Figure 11 **a**) deviates from describing a regular flat polygon. These descriptors build up on ideas from piecewise-linear (PL) topology. A detailed introduction to this field can be gained from [51]. A short review of relevant notions is presented below.

The setting that is being discussed here is that of a point cloud $S \subset \mathbb{R}^3$ lying on or close to a 2D surface. The shape of that surface can be approximated using a triangulation $T$ that is computed from $S$. More specifically, $T$ is a complex that is built from vertices, edges, and triangles that only intersect in building blocks from $T$, namely in common “faces”: triangles intersect in edges or vertices, edges intersect in vertices. The underlying structure of such a $T$ is also referred to as a simplicial complex; its building blocks are “simplices”.

The coarsest shape approximation of $S$ is its convex hull, which, in 3D, can be built from 3-dimensional simplices (tetrahedra). Since our focus lies on 2D structures, this case is to be avoided, which can be achieved in different ways. The one that is presented here is based on the assumption that the sampled surface is tame enough so that sufficiently small data patches can be easily projected onto the plane. Therefore, we shift the discussion towards point clouds $S \subset \mathbb{R}^2$. In our setting, the topological information (What data points are connected via edges / triangles?) gained from triangulations of data patches in the plane can be transferred back to the corresponding patches of the original point cloud in 3D.

The shape of $S \subset \mathbb{R}^2$ can be approximated using a family of nested complexes, the so-called $\alpha$-complexes [52, 53, 18], which highlight varying degrees of connectedness of that shape (connected components, holes). The parameter $\alpha^2$ is an upper bound on the radii of circumcircles of the building blocks of a special triangulation of the convex hull of $S$, the Delaunay triangulation [54]. Thus, $\alpha$-complexes can be helpful in representing the data's shape more accurately, for example in the case of holes in virus capsids. Note that while distance information may be distorted when projecting onto the plane, thus also having an effect on the $\alpha$ parameter, this approach still yields faithful local information within the assumption of “tame” 2D surfaces.

In the case of a triangulated 2D manifold without boundary, the so-called star of a given vertex consists of all triangles (and their edges and vertices) that have this vertex in common, see for example Supplementary Figure 11 **a**. More generally, the star consists of all simplices of $T$ that touch on the vertex in question, as well as all of their faces. The link of a vertex consists of all simplices of the vertex's star that don't contain the vertex at hand. In the example of a star made up of triangles, this translates into edges and vertices not touching the query vertex.

Within the computation of the PL descriptor, every particle assembling on a 2D structure will be considered a vertex in a triangulation. Focusing on every vertex's star, the associated features describe the star's geometry: the central angles (i.e. angles incident to the query vertex), every triangle's circumcircle radius, as well as the radii of their inscribed circles, and their surface areas. These geometric features are computed for the 3D information that is gained when mapping the topological information from the 2D triangulation back to the original 3D point cloud.

#### 2.2. $\alpha$-complex descriptor

Similar to the PL descriptor, this one focuses on stars of vertices, yet in this case, the stars can reflect incompleteness of data based on the $\alpha$ parameter (see Supplementary Figure 11 **b**). Besides geometric features describing angles and surface areas, this descriptor supports the analysis of link topology and link architecture. Under the tameness assumption of triangulated

2D surfaces, a link's dimension is at most 1. For a given link $L$ of dimension 1, the features that are of special interest are the Euler characteristic $\chi(L)$ and a simplicial isomorphism invariant $e(L)$ describing the type of link [55]:

$$\chi(L) = (\text{number of vertices in } L) - (\text{number of edges in } L) \tag{14}$$

$$e(L) = 1 - \frac{\text{number of vertices in } L}{2} + \frac{\text{number of edges in } L}{3}. \tag{15}$$

These allow us to judge whether the support shows any characteristic of e.g. pentagonal or hexagonal arrangement, or whether the neighborhood is complete in this sense. This makes the $\alpha$-complex descriptor suitable for analysis of (semi-) regular arrangements, such as lattices.

#### 2.3. SHOT descriptor

SHOT (signature of histograms of orientations) is suited to discriminate between particles based on the spatial arrangement of their nearest neighbors. For rotational invariance, every spherical support is rotated so that the query particle is in its canonical orientation. Then, the spherical support is subdivided into cones, which are further subdivided into radial shells (Supplementary Figure 11 **c**). Based on this, the SHOT descriptor associates to every particle a histogram describing how the individual bins are occupied. Comparison of these histograms can uncover common positional patterns and highlight regions for further investigation, such as an analysis on whether some positional pattern translates into an orientational one.

| Use case | Support | Axis | Radius (nm) | Height (nm) | AD (rad) | RDZ (rad) | Relevant physiological data | Section |
|---|---|---|---|---|---|---|---|---|
| **NPC** | cylinder | $z$ | 18.4 | 6.1 | 0.7 – 0.9 | 0.5 | 66.6 nm mean diameter [6], expected AD between neighboring SUs $\frac{1}{4}\pi \approx 0.79$ rad | 3.1 |
| **VLP** | cylinder | $-z$ | 15.2 | 1 | | 0.1 | 6.8 nm mean NN distance | 3.1 |
| | sphere | | 15.2 | | | | | 3.2 |
| **MT** | cylinder | $x+y$ | 7 | 2.3 | | | 4.5 nm mean NN distance | 3.1 |
| **Capsid** | sphere | | 10.8 | | | | 8.9 nm mean NN distance | 3.2 |
| **Synth** | sphere | | 40.8 | | | | 10.2 nm mean NN distance | 3.3 |
| | cylinder | $y$ | 9.8 | 12.2 | | | | |
| | cone | $z$ | 19.6 | 40.8 | | | | |
| **Ribo** | sphere | | 9 | | | | 13.1 nm distance center to entry, 10.5 nm distance center to exit | 3.3 |

**Supplementary Table 1**
Parameters used in TANGO for use cases. Abbreviations: AD = angular distance, RDZ = rotational deviation from $z$-axis = $|\zeta_z - \|\zeta\||$, NPC = nuclear pore complex, VLP = virus-like particle, MT = microtubules, SU = subunit, Synth = synthetic data, Ribo = ribosomes, NN = nearest neighbor. The parameters for the support in the CR case were chosen as 30 voxels for height, 10 voxels for radius. For VLP, MT, and the capsids, linear combinations of median of nearest neighbor distances and standard deviation of nearest neighbor distances were chosen after sampling some options of combinations that yielded supports containing the first shell around each query particle. In the case of ribosome data, the sphere radius was chosen to remain consistent with the original study [4]. In the cases of MTs and VLPs, support radii were chosen to obtain mostly complete first shells.

| Parameter name (used in novaSTA) | Value - one per iteration unless it was constant |
|---|---|
| coneAngle | 80 80 80 40 40 40 20 20 20 20 |
| coneSampling | 10 10 10 8 8 8 5 5 5 5 |
| inplaneAngle | 80 80 80 40 40 40 20 20 20 20 |
| inplaneSampling | 10 10 10 8 8 8 5 5 5 5 |
| lowPass | 10 10 10 10 10 12 12 12 12 14 |
| highPass | 1 |
| symmetry | 1 |
| PixelSize | 4.752 |
| BoxSize | 52 |

**Supplementary Table 2**
Subtomogram averaging parameters use for alignment of microtubules from Section 3.1.

| $C_6$ vs. • | $p$-value | Mann-Whitney $U$ statistics | sample size |
|---|---|---|---|
| $C_2$ | $8.4 \cdot 10^{-19}$ | $2.7 \cdot 10^5$ | 844 |
| $C_3$ | $4.0 \cdot 10^{-20}$ | $2.6 \cdot 10^5$ | 844 |
| $C_4$ | $3.6 \cdot 10^{-35}$ | $2.3 \cdot 10^5$ | 844 |
| $C_5$ | $8.6 \cdot 10^{-51}$ | $2.0 \cdot 10^5$ | 844 |
| $C_7$ | $1.5 \cdot 10^{-49}$ | $5.1 \cdot 10^5$ | 844 |
| $C_8$ | $4.5 \cdot 10^{-46}$ | $5.0 \cdot 10^5$ | 844 |
| $C_9$ | $7.6 \cdot 10^{-47}$ | $5.0 \cdot 10^5$ | 844 |
| $C_{10}$ | $9.5 \cdot 10^{-46}$ | $5.0 \cdot 10^5$ | 844 |

**Supplementary Table 3**
Results of Mann-Whitney $U$ test, comparing data collected under the assumption of $C_6$-symmetry with data collected under the assumption of $C_n$-symmetry.

| $C_n$ | min | Q1 | median | Q3 | max |
|---|---|---|---|---|---|
| $C_2$ | 0.03 | 0.32 | 0.45 | 0.90 | 1.00 |
| $C_3$ | 0.00 | 0.11 | 0.55 | 0.90 | 1.00 |
| $C_4$ | 0.01 | 0.28 | 0.50 | 0.81 | 1.00 |
| $C_5$ | 0.00 | 0.24 | 0.50 | 0.75 | 1.00 |
| $C_6$ | 0.00 | 0.59 | 0.79 | 0.91 | 1.00 |
| $C_7$ | 0.00 | 0.25 | 0.52 | 0.77 | 1.00 |
| $C_8$ | 0.00 | 0.29 | 0.51 | 0.77 | 0.99 |
| $C_9$ | 0.00 | 0.21 | 0.53 | 0.79 | 1.00 |
| $C_{10}$ | 0.00 | 0.28 | 0.52 | 0.78 | 1.00 |

**Supplementary Table 4**
Quantiles corresponding to angular score analysis under the assumption of $C_n$-symmetry.

| Parameter name (used in novaSTA) | Value - one per iteration unless it was constant |
|---|---|
| coneAngle | 30 30 30 20 20 20 10 10 10 10 10 10 10 10 10 |
| coneSampling | 5 5 5 4 4 4 2 2 2 2 2 2 2 2 2 |
| inplaneAngle | 80 60 60 45 45 30 30 20 20 20 10 10 10 10 10 |
| inplaneSampling | 7 5 5 4 4 3 3 2 2 2 2 2 2 2 2 |
| lowPass | 16 16 16 16 16 18 18 18 18 18 18 18 18 18 18 |
| highPass | 1 |
| symmetry | 1 |
| PixelSize | 5.316 |
| BoxSize | 72 |

**Supplementary Table 5**
Subtomogram averaging parameters use for alignment of VLPs from Section 3.2.2.

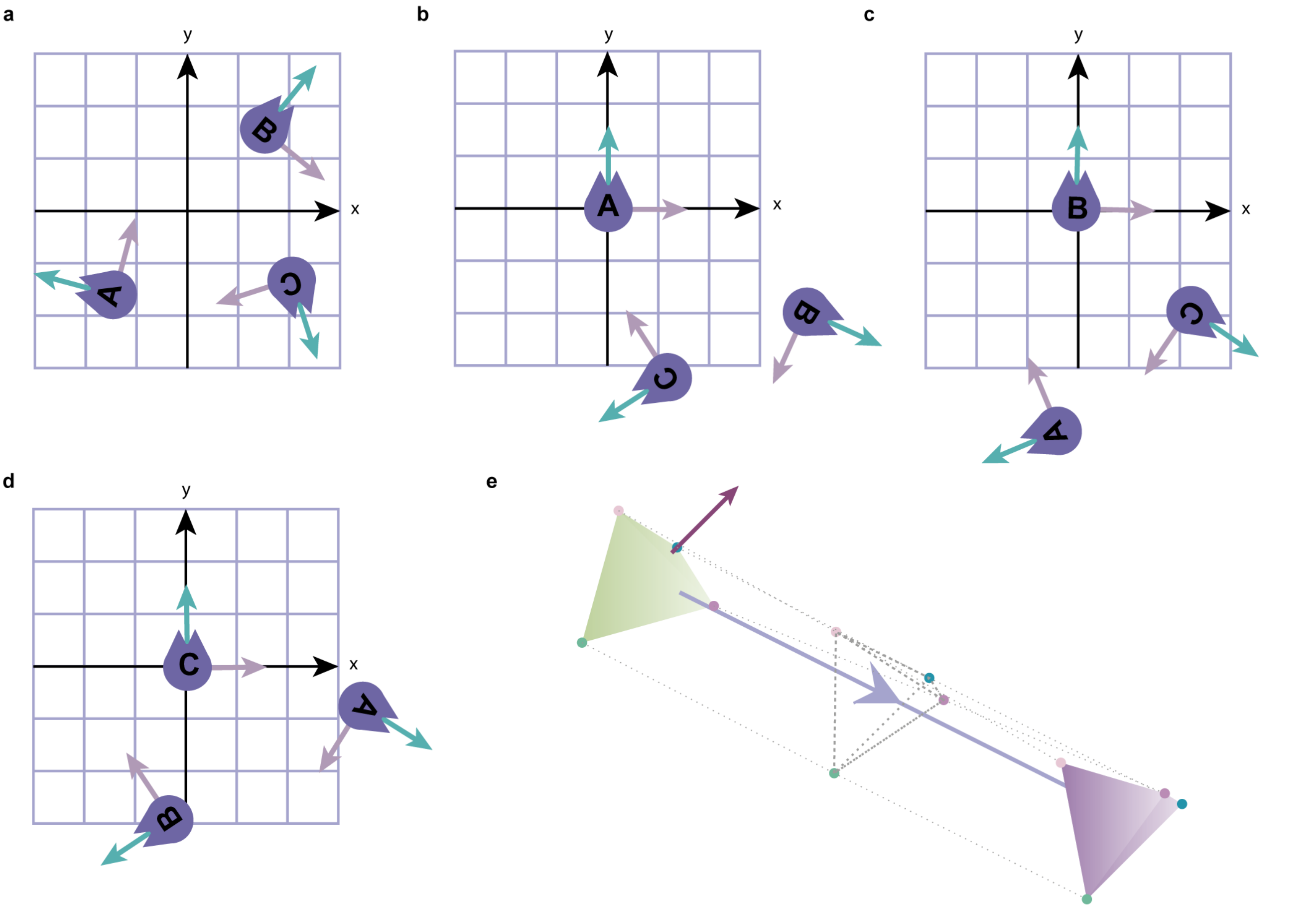


**Supplementary Figure 1: a**. Initial configuration of planar cats A, B, C with indicated orientations. **b**. Planar cat A aligned with canonical orientation and shifted to origin. The other cats are shifted and rotated accordingly, while respecting relative orientaions and distances with respect to cat A. **c, d**. Same as in **b**, but for planar cats B and C, respectively. **e**. Visualization of a rigid motion. This motion in 3D acts on the left tetrahedron by rotating it around the purple axis, and translating it along the light lavender translation vector connecting it to the tetrahedron on the right. The latter is the final position and orientation of the former.

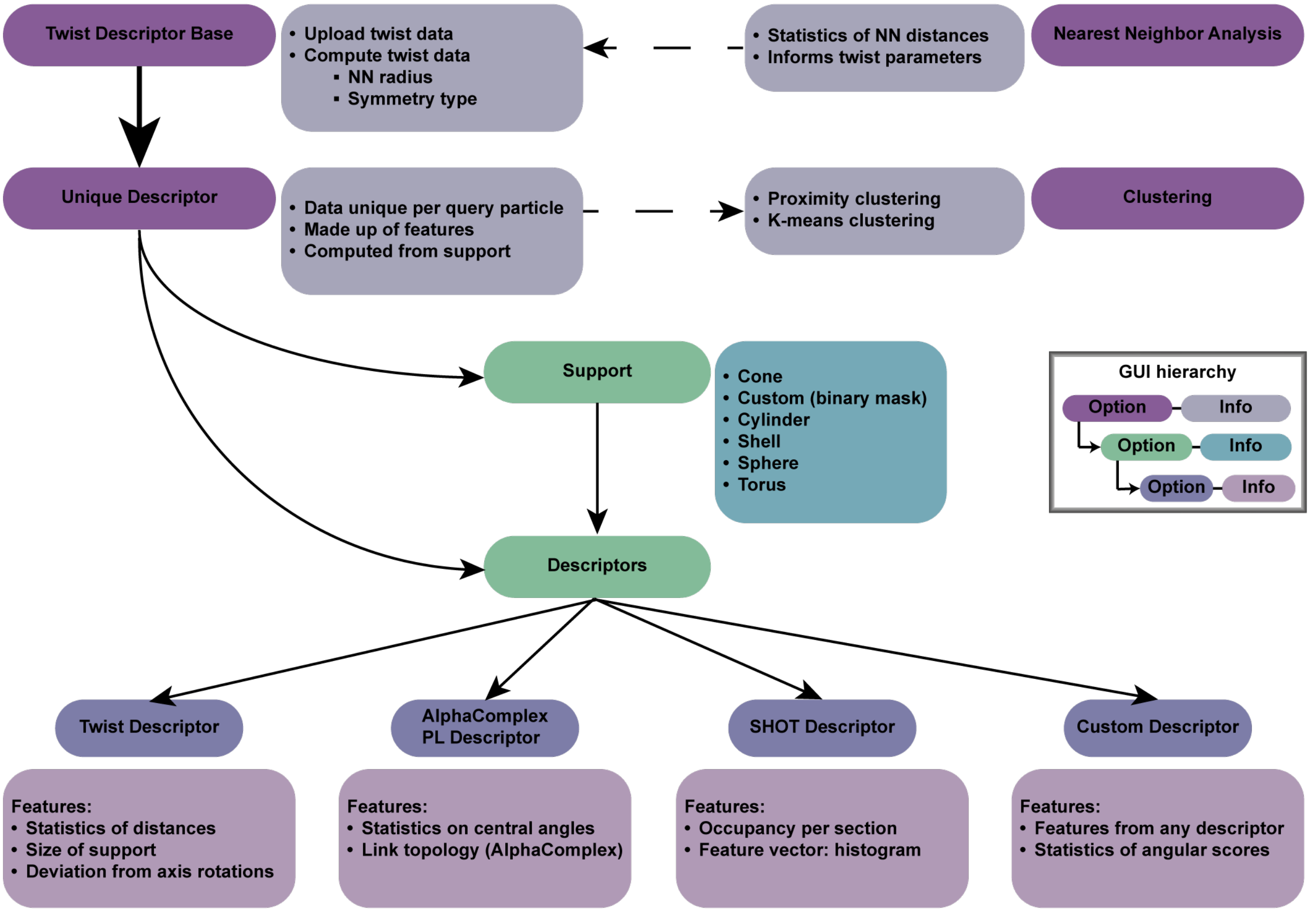


**Supplementary Figure 2:** A diagram explaining the structure of the TANGO graphical user interface (GUI). Dashed arrows indicate the order as proposed by the framework. The thick arrow indicates that the "Twist Descriptor Base" is a prerequisite for the computation of any "Unique Descriptor".

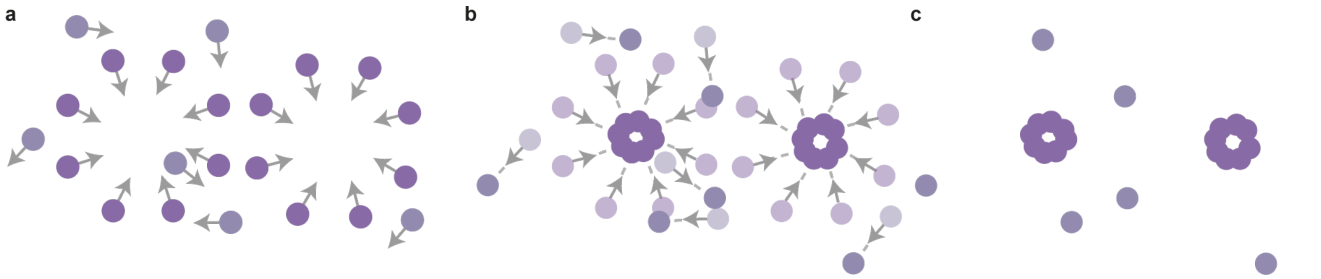


**Supplementary Figure 3:** Visual explanation of the preprocessing step used for cleaning and affiliation of the NPC CR. **a**. Initial state of particles. Gray arrows indicate the intrinsic $-x$ axis. Purple particles denote asymmetric units of the CR (true positives), while lavender particles represent false positives. **b**. Shift of particles along the $-x$ axis. Starting positions are shown in lighter hues than ending positions. **c**. Final positions of shifted particles from panel **a**. Shifting brings the true positives closer together, whereas the false positives move randomly.

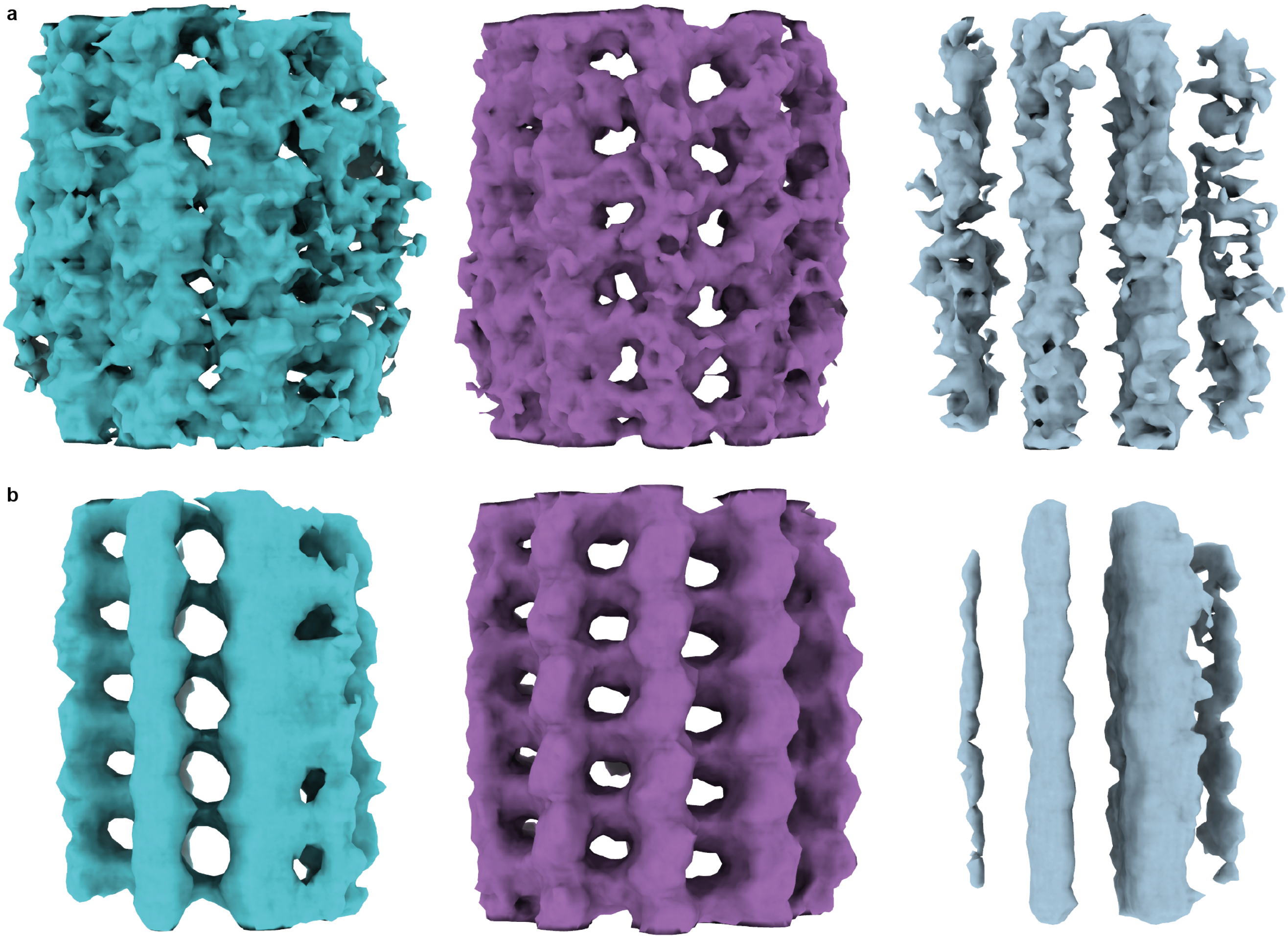


**Supplementary Figure 4:** Subtomogram averaging of microtubules from Section 3.1. **a**. Left: Initial average of all particles (true and false positives) from TM. Right: Average after 10 STA iterations. **b**-**c**. Same as in **a**, but for the cleaned list and the removed-particle list, respectively. All initial averages resembled the template to some extent. With further alignment, the combined and removed sets degraded, while the cleaned set improved, supporting the effectiveness of the cleaning. Some resemblance to microtubules remained in both combined and removed sets—expected in the former due to true particles, and in the latter possibly reflecting a small number of false negatives.

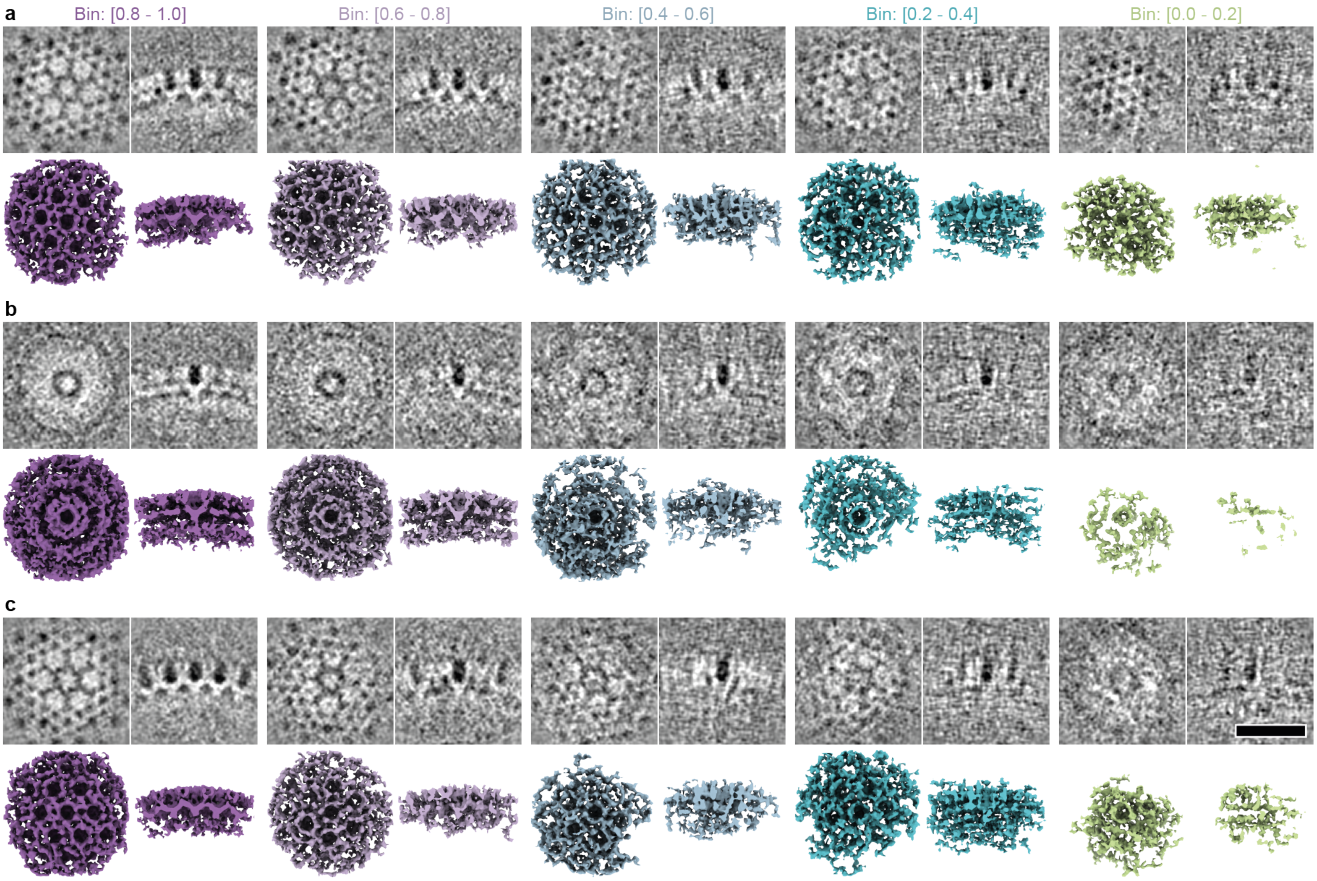


**Supplementary Figure 5:** Subtomogram averaging of CA-SP1 capsid from VLPs of immature HIV-1. **a**. STA maps created by averaging particles with different ranges of their mean angular scores. Each map is represented by 2D slices showing its top view (top left), its side view (top right) and by corresponding top (bottom left) and side views (bottom right) of its iso-surface representation. **b**. Initial STA averages created by averaging particles after randomizing their in-plane angles. Apart from the randomization, the particle lists were identical as those in **a**. **c**. STA maps corresponding to the initial averages from **b** after 15 iterations of STA alignment. Scale bar 20 nm.

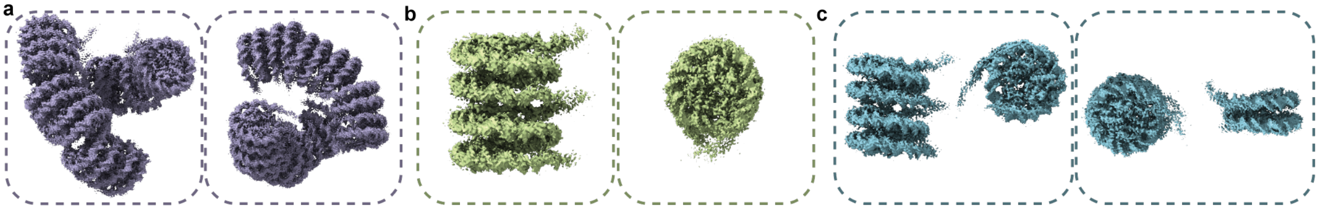


**Supplementary Figure 6:** **a**. Visual representation of the particle list deduced from EMD-2601 representing the helical arrangement. Left: side view. Right: top view. **b**. Visual representation of the particle list deduced from EMD-13365 representing the stacked nucleosomes. Left: side view. Right: top view. **c**. Visual representation of particle list deduced from EMD-13363 representing the trinucleosome. Top: side view. Bottom: top view.

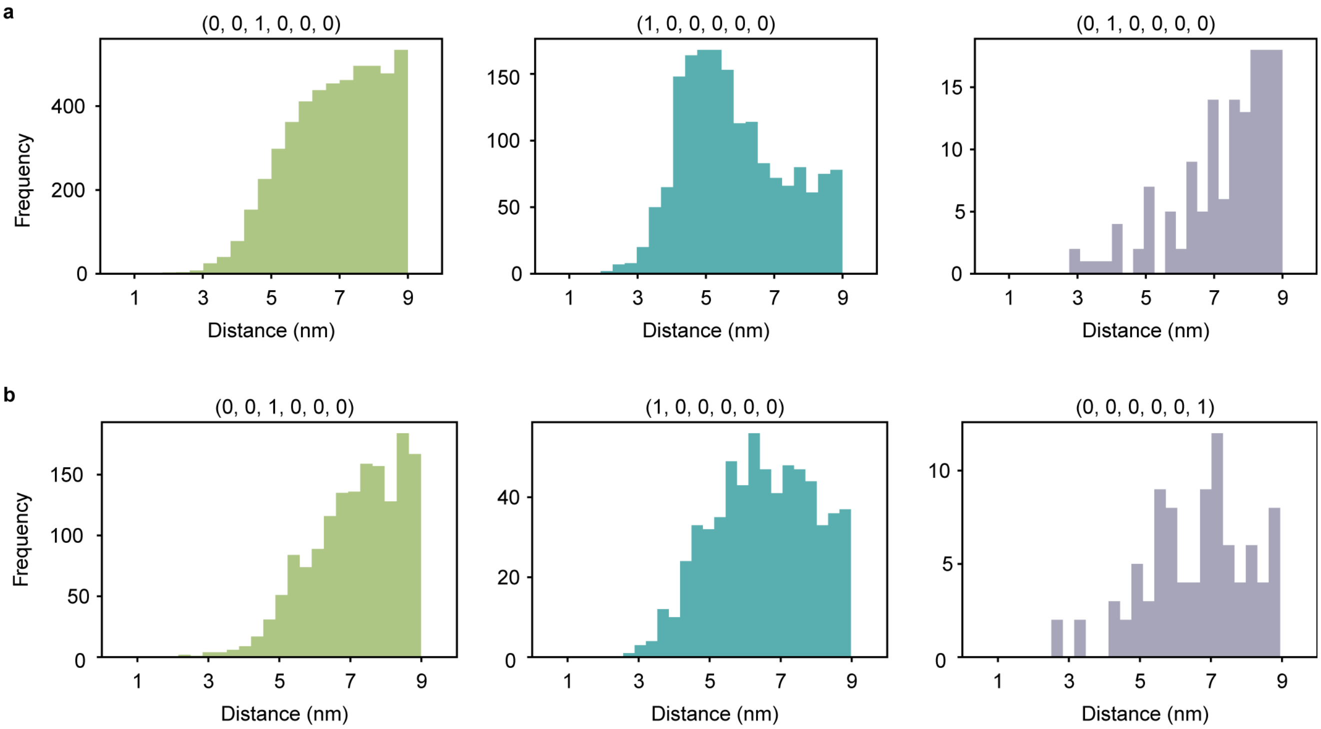


**Supplementary Figure 7:** Distance distributions per SHOT footprint for ribosome analysis. SHOT footprints within the given SHOT parameters (number of cones: 6, number of shells: 1) are shown above every histogram. The entries of the depicted footprints are to be read as follows: The first entry yields the number of neighboring particles in the cone around the query particle's intrinsic $z$-axis, the second entry does the same for the intrinsic $x$-axis, and so on continuing with the intrinsic $y, -z, -y, -x$-axes in that order. **a**. Distance distributions in the untreated case. **b**. Distance distributions in the treated case.

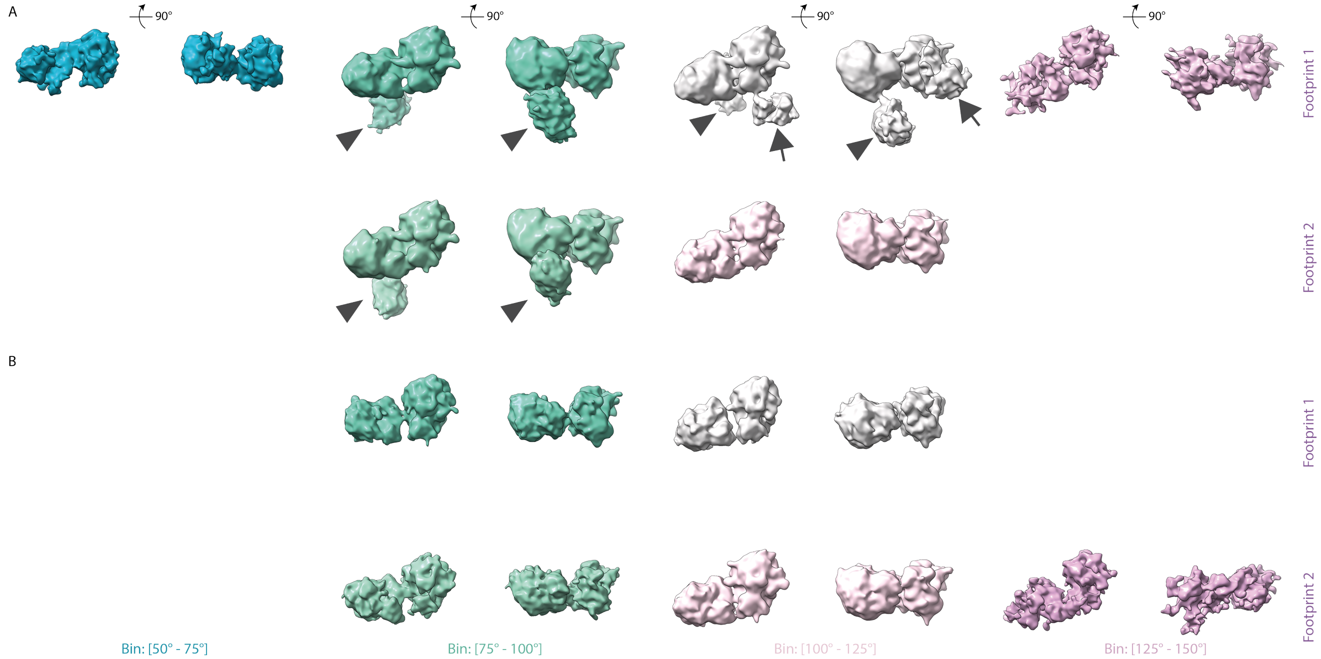


**Supplementary Figure 8:** **a**. Selection of STA results from the detected particles in the untreated case. The relevant SHOT footprints (FP) and bins for different angular distances are mentioned. Rotations are by 90° as indicated for the first pairs. The arrowheads point to a third ribosome that emerged during averaging, suggesting a pattern of three ribosomes with relatively preserved mutual orientation. The arrows point to a fourth ribosome present in the average. **b**. Same as in **a** for the treated condition. Note the lack of any additional densities apart from the pairs.

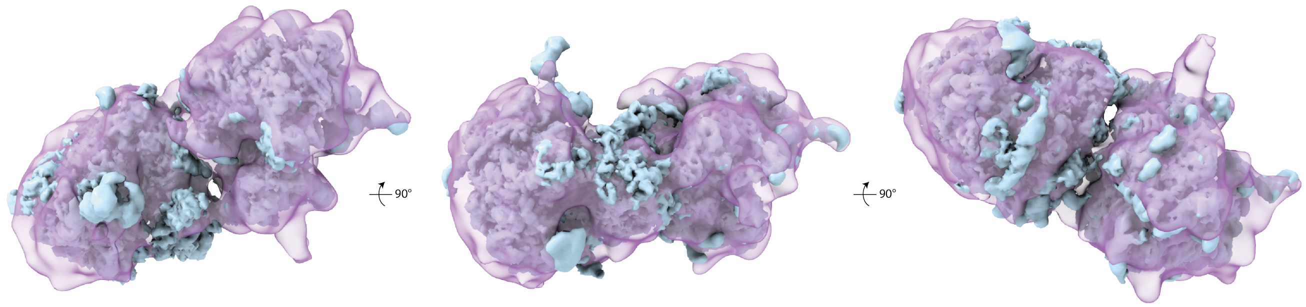


**Supplementary Figure 9:** The map of the stalling disome (EMDB-10398) overlayed with the STA map of the pairs from the most occupied bin of the third footprint of the treated condition.

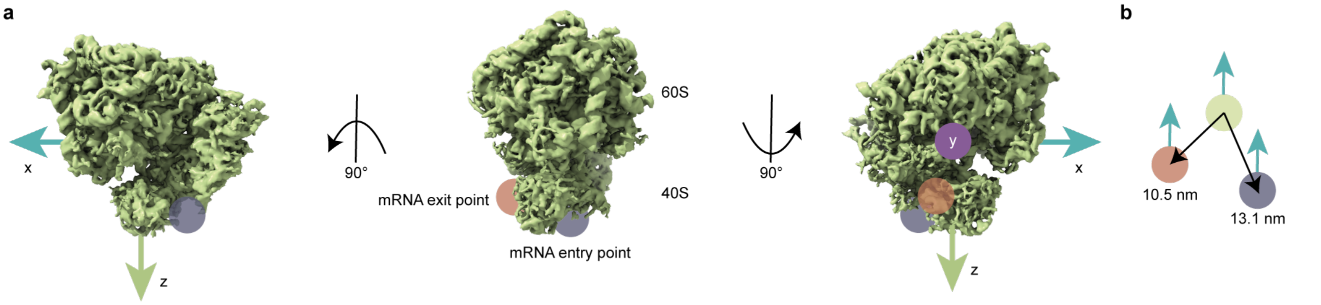


**Supplementary Figure 10:** **a**. Model of a ribosome with indicated entry and exit points for mRNA. Rotated ribosomes include indicated normal vectors corresponding to the reference's orientation. **b**. Distances to center are shown in with colors corresponding to entry and exit points as in **a**. When shifting the particle list containing positions of ribosome centers to entry and exit points, orientations are kept the same as indicated by the turquoise arrows. Compare [4], Fig. S12.

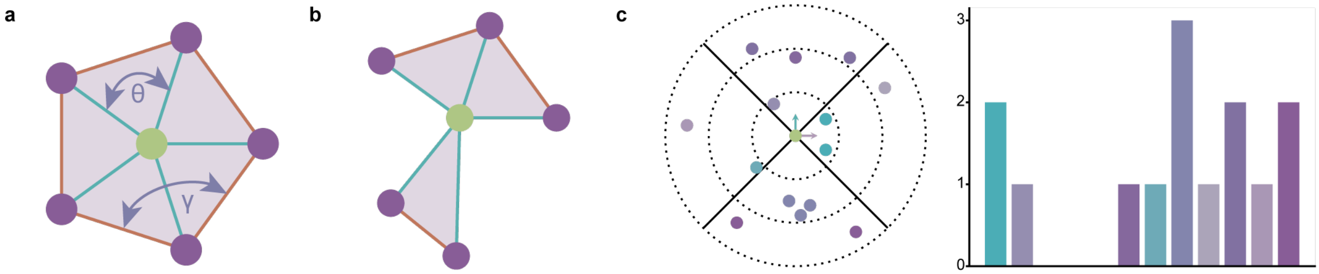


**Supplementary Figure 11:** **a**. The star of a vertex with indicated central angle $\theta$ and internal angle $\gamma$. **b**. A star can also reflect holes in the data, resulting for example in a link with two connected components (orange). **c**. SHOT computes histograms describing the occupancy of individual bins of a subdivided spherical support. A color-coded histogram corresponding to the subdivided support is shown on the right. The SHOT parameters in this example are: number of cones = 4, number of shells = 3.
